# Indexcov: fast coverage quality control for whole-genome sequencing

**DOI:** 10.1101/148296

**Authors:** Brent S. Pedersen, Ryan L. Collins, Michael E. Talkowski, Aaron R. Quinlan

## Abstract

The BAM^1^ and CRAM^2^ formats provide a supplementary linear index that facilitates rapid access to sequence alignments in arbitrary genomic regions. Comparing consecutive entries in a BAM or CRAM index allows one to infer the number of alignment records per genomic region for use as an effective proxy of sequence depth in each genomic region. Based on these properties, we have developed *indexcov*, an efficient estimator of whole-genome sequencing coverage to rapidly identify samples with aberrant coverage profiles, reveal large scale chromosomal anomalies, recognize potential batch effects, and infer the sex of a sample. *Indexcov* is available at: https://github.com/brentp/goleft under the MIT license.

## Introduction

Whole genome sequencing (WGS) studies produce massive datasets that cost thousands of dollars per sample and often require hundreds or thousands of hours to analyze with intense computational requirements. While verifying the integrity and quality of the resulting sequence data is crucial, it remains difficult owing to the size of the data. For example, a single aligned BAM file from 30X WGS typically results in hundreds of millions of alignment records, requiring at least 100 gigabytes of storage in BAM format. The simple act of iterating through every alignment record (without any analysis or computation) can consume hours of processing time. Assessing the depth and breadth of DNA sequence coverage in a WGS sample is a necessary precursor to variant discovery, as depth of coverage drives the power to detect genetic variation, especially in the case of heterozygous sites^2,3^. Coverage information is critical when detecting copy number (CNV) and structural variation (SV), as greater sequencing depth increases power to detect smaller CNVs and increases the probability that an SV breakpoint will be captured by multiple, independent sequence fragments^4,5^. However, existing quality-control (QC) tools^1,6,7^ do not provide rapid visualizations of genome-wide or targeted estimates of sequence coverage for multiple samples, which if aberrant, can confound downstream analyses. As a result, when studying large cohorts, a problematic sample may remain undetected until after these steps are completed. Therefore, it is critical to assess coverage profiles in a cohort as early as possible to identify problematic samples before proceeding with further analyses.

In addition to single-sample problems such as missing data, it is important to look for batch effects and systematic artefacts in sequencing projects^8^. The ability to rapidly summarize coverage across all samples after sequencing and alignment can identify problematic samples that require additional sequencing, or that should be excluded from subsequent analysis. In an effort to address the quality control needs of WGS studies, we introduce *indexcov* as a new software package to quickly estimate the depth and consistency of sequence coverage in a BAM or CRAM file. *Indexcov* interrogates the entire genome of a sequenced sample using either a linear BAM index (default resolution: 16,384bp) or a CRAM index (variable default resolution) to generate rapid estimates of coverage depth at every position on every chromosome. Using this efficient approach, *indexcov* is able to infer sample sex, perform a principal components analysis to identify batch effects, and reveal coverage anomalies much more quickly (~seconds per genome) than existing methods. In addition, *indexcov* produces interactive, web-based plots that permit users to visualize and investigate the coverage profiles and related QC metrics for each sample, both in a targeted fashion (e.g. individual chromosomes) or summarized across the whole genome.

## Results

### Estimating WGS Sequence Depth from Alignment Indexes

The BAM index enables random access in a coordinate-sorted BAM file, facilitating rapid interrogation of sequence reads aligned to arbitrary genomic regions in a reference genome. It provides a linear index that saves the file and block offset of the first alignment to start in each consecutive 16,384bp “tile” of the genome. *Indexcov* iterates over all tiles per chromosome, recording the number of bytes that exist in each tile; the median of these proxy coverage values is used to establish a baseline coverage level for the average tile per chromosome. For example, imagine the median of all genome-wide tiles consumes ~32 kilobytes. If we then identify a large stretch of adjacent tiles that consume ~16 kilobytes (i.e., half the median), this may be evidence of a heterozygous deletion in a diploid organism. While the CRAM index does not have fixed-width tiles in terms of genomic bases, it can be used in a similar fashion. Since each sample will have different regions for each container (chunk) in the CRAM index, *indexcov* divides the CRAM chunks into 16KB bins to normalize across samples. This results in a loss of resolution, but this approach still enables the detection of large coverage anomalies.

### Comparison of Coverage Inferred by *Indexcov* to Empirical per-Base Sequencing Depth

A potential complication with the *indexcov*’s coverage estimation approach is that individual tiles may differ from the median value in a number of ways that are not due to bona *fide* changes in the depth of coverage or DNA dosage in that sample. For example, tiles with apparently high coverage may reflect situations where there are many split reads, which have more SAM tags, thereby increasing the number of bytes required by each alignment. Nonetheless, we found that the coverage estimated by *indexcov* is well correlated with the actual depth calculated by aggregating per-base depth calls from the samtools^1^ “depth” command into 16,384bp tiles (**Figure 1**). We conducted this analysis on chromosome 1 for a single human BAM file aligned to reference assembly GRCh37 (NA12878, available here: ftp://ftp.sra.ebi.ac.uk/vol1/ERA172/ERA172924/bam/NA12878_S1.bam), and, for the purpose of comparing the depth reported by samtools to the scaled value in *indexcov*, we divided the depth from samtools in each window by the overall median.

**Figure 1.**
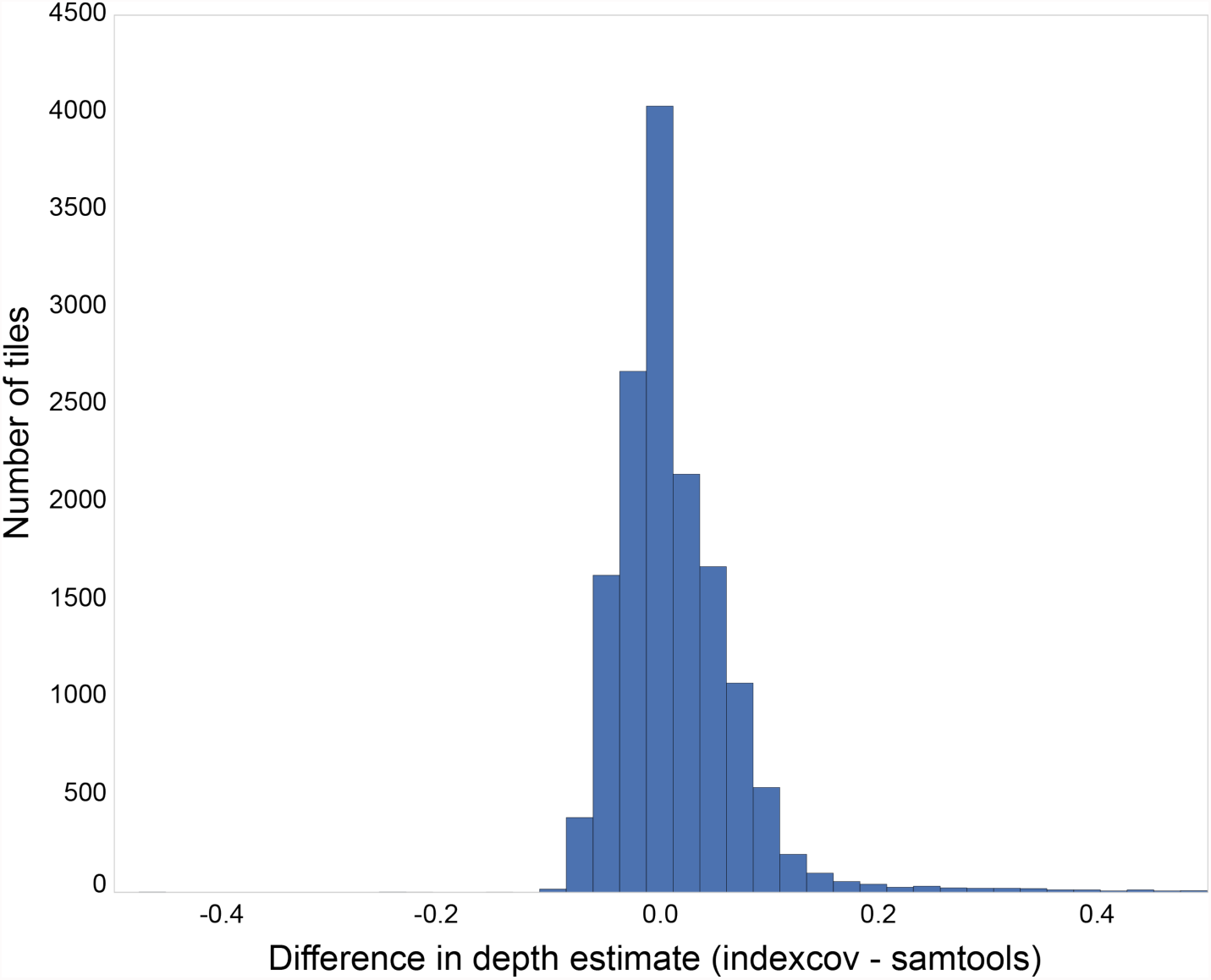
Difference between median-scaled sequencing depth in 16,384bp bins from samtools, which recovers per-base depth from the BAM file, and *indexcov*, which estimates coverage from the BAM index. Samtools required ~8 minutes to compute depth in 16.4kb bins for chr1, whereas indexcov estimated depth in 16.4kb bins genome-wide in under one second. Pictured here is a summary from NA12878 chromosome 1: x-axis values indicate the relative difference in normalized coverage estimates between samtools and *indexcov* in 16.4kb bins.

Whereas this analysis required ~8 minutes of CPU time for samtools on chromosome 1, *indexcov* completed the analysis in 1 second for the entire genome. Indeed, less than 3% of the 15,196 tiles (each 16,384bp in size) on chromosome 1 differed between *indexcov* and samtools by more than 5% of the normalized coverage values. Overall, the strong correspondence between normalized coverage values from indexcov and samtools suggests that indexcov is an effective--and vastly more computationally efficient-proxy for estimating large-scale coverage values across an entire aligned WGS sample stored in BAM or CRAM format.

### Analysis of Genome-Wide Coverage with *Indexcov*

The most intuitive output created by *indexcov* is an estimate of genome-wide coverage, which can be run on individual samples or simultaneously across large batches of samples for cohort-wide QC and coverage analyses. *Indexcov* produces one coverage plot for each chromosome via an interactive plot contained in an HTML file (details below), as well as a BED file containing the scaled coverage values for each sample at each 16KB tile, thereby enabling custom visualizations if desired. An example coverage plot is shown in **Figure 2A** for chromosome 15, reflecting coverage profiles for 45 human WGS samples from an ongoing study at the University of Utah. Since chromosome 15 is acrocentric and its centromere is N-masked in the human reference genome assembly, we see no reads aligned to the first ~20Mb (far left) of the plot. Aside from a highly repetitive region downstream of the centromere, most of the coverage values for the majority of samples are centered at a scaled coverage of 1, which corresponds to a diploid copy state for chromosome 15. However, a single sample (highlighted in green) has a scaled coverage value of ~0.5 across a 10 megabase region (23-33Mb), which is consistent with a large deletion that resulted in a genetic diagnosis of Angelman syndrome for this individual. While *indexcov* is not intended as a general purpose CNV detection tool, it serves as an effective method for visual identification of large anomalies such as this Angelman Syndrome deletion. **Figure 2B** shows this same coverage information as a reverse cumulative density function (CDF). Like the first coverage plot, this view also clearly highlights the aberrant sample with the Angelman Syndrome deletion, as evinced by ~10% of chr15 being covered at a lower scaled coverage value than the other 44 samples. Most samples have a steep slope at a scaled coverage value of ~1, reflecting the fact that the majority of genome tiles for these samples are very close to a scaled coverage of 1. However, when a sample has much greater variability in scaled coverage (e.g., sample in red), the slope when passing through a scaled coverage of 1 will be far less steep. Chromosomes with high GC content (such as 19, 22, 17,16 in humans) will vary more widely in slope consistent with GC-correlated biases in sequencing depth introduced by PCR^9^. 

**Figure 2.**
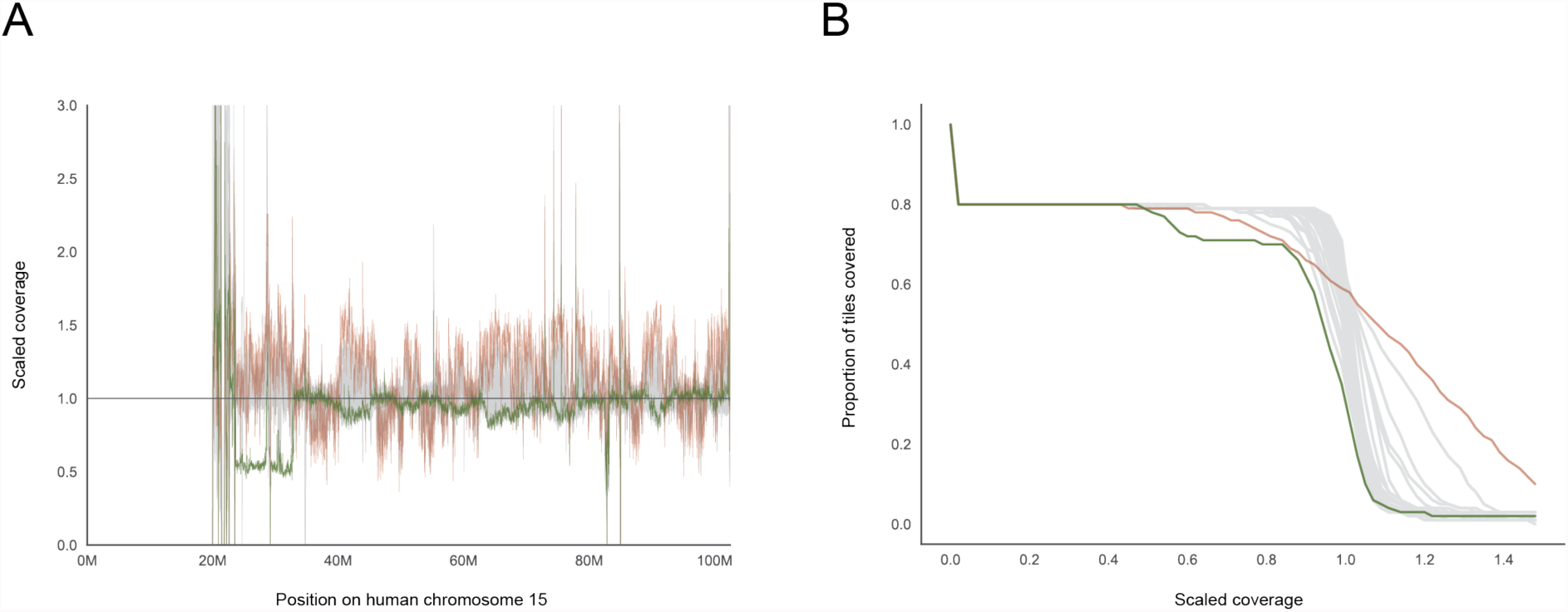
Coverage profiles for 45 human WGS samples on chromosome 15. The estimated coverage along the chromosome is shown in (**A**) and an alternative representation showing the proportion of tiles covered at a certain depth and as the lower path in (**B**). The sample highlighted with a green line has a ~1 0MB deletion just after the (acrocentric) centromere that has been previously associated with Angelman syndrome. The crimson line tracks a sample with a large variability in coverage; samples like this one will have many spurious CNV calls. These plots are interactive in the *indexcov* output, allowing users to hover and identify samples of interest.

### Sex Inference

Sequencing depth is an especially effective metric for ploidy and sex inference since human males typically have only one X chromosome and human females lack Y chromosomes. Using these expectations, and as a demonstration of the utility of *indexcov* to rapidly facilitate useful cohort-wide QC in WGS studies, we used *indexcov* to infer the ploidy of the sex chromosomes in a cohort of 2,076 from 519 ‘quartets’ (proband, unaffected sibling, and two parents) samples as part of a recent analysis of autism spectrum disorder (ASD) simplex families^10^ (**Figure 3**). Most samples cluster as either male (XY genotype; X=1, Y=1) or female (XX genotype; X=2, Y=0). However, five samples cluster in two non-canonical locations, one at (XYY inferred genotype; X=1, Y=2) and a second cluster at (inferred genotype XXY; X=2, Y=1). These non-canonical clusters of samples indicate the presence of rare supernumerary sex aneuploidies, some of which have been previously suggested as pathogenic in ASD^11^. Notably, although these *indexcov* analyses were performed blind to all prior genetic knowledge for these samples, all of the sex aneuploidies discovered by *indexcov* had been previously discovered by an earlier analysis of these samples using SNP microarray^12^, thereby demonstrating that *indexcov* can accurately detect sex chromosome anomalies also corroborated by preexisting methods. Finally, a single sample appears just below X=0, Y=0, which was the result of a truncated BAM index. While this sample issue was easily resolved by re-indexing the original BAM file, it serves as a valuable example of technical problems that can be identified by *indexcov*. While *indexcov* natively assumes that the sex chromosomes are X and Y, the user can also override these defaults when necessary, such as for non-human organisms or alternative reference assemblies.

**Figure 3.**
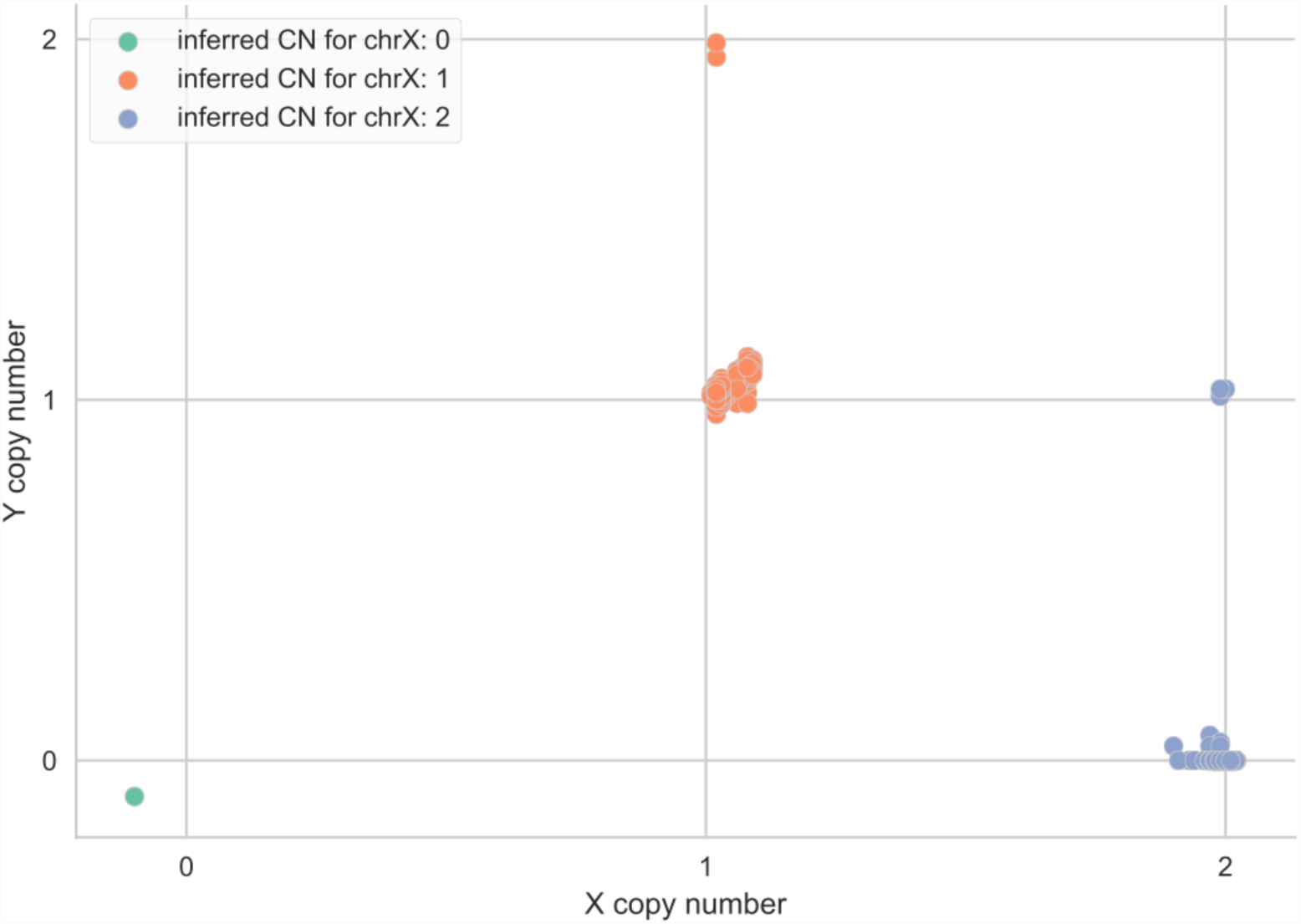
Sex inference plot for a cohort of 2,076 human WGS samples analyzed with *indexcov*. Samples projected on this plot represent ~30-40X human WGS from 519 “quartet” families recently analyzed as a study of simplex autism (Werling et al., 2017). The x-axis shows the copy-number for chrX and the Y-axis shows the copy-number for chrY inferred by *indexcov*. Sex is inferred from the copy-number of X. As expected, we see two dominant clusters of samples, one of males (X=1 and Y=1) and one of females (X=2 and Y=0). Notably, *indexcov* further identifies samples with supernumerary sex chromosome aneuploidies (XXY and XYY), which had previously been identified by SNP microarray analysis (Sanders et al., 2015). The green point in the lower left just below the origin represents a sample with no apparent coverage on chromosomes X or Y due to a truncated BAM index file, which can be rapidly corrected once identified by *indexcov* QC.

### Sequencing Batch Effect Detection with Principal Component Analysis

*Indexcov* uses **Principal** Component Analysis (PCA) to identify batch effects or other major discrepancies among groups of WGS samples. As *indexcov* creates the coverage plots for each chromosome, it simultaneously appends a large array for each sample that contains the scaled coverage values for the entire genome. Once all chromosomes are completed, *indexcov* performs PCA on the scaled coverage values and projects all samples onto the first 5 principal components, finally outputting two PCA plots: the first and second principal components, and the first and third principal components. PCA visualization enables the detection of fundamental differences in sets of samples, such as WGS samples that were sequenced with or without PCR amplification (**Supplemental Figure 1**).

### Identifying Aberrant Genome-Wide Coverage Profiles

While calculating scaled coverage for each chromosome, *indexcov* tallies several other informative metrics. First, it measures the proportion of 16kb tiles that had a scaled coverage between 0.85 and 1.15, as well as scaled coverage <0.15, or >1.15. In our experience, these simple cutoffs work well to differentiate samples with highly aberrant coverage anomalies from normal, uniformly covered samples, although we have also found that the results are quite stable even when the cutoffs are changed moderately (data not shown). The resulting “tile plots” convey the proportion of low values (< 0.15) versus the proportion of tiles with values outside of 0.85 - 1.15. An example application of this approach is illustrated in **Figure 4** for the same cohort of 2,076 WGS samples show in **Figure 3**. This method highlights a single sample with a very large value on the X-axis indicating that it is missing data for many tiles. In fact, this plot led us to realize that the BAM index for that sample had been truncated.

**Figure 4.**
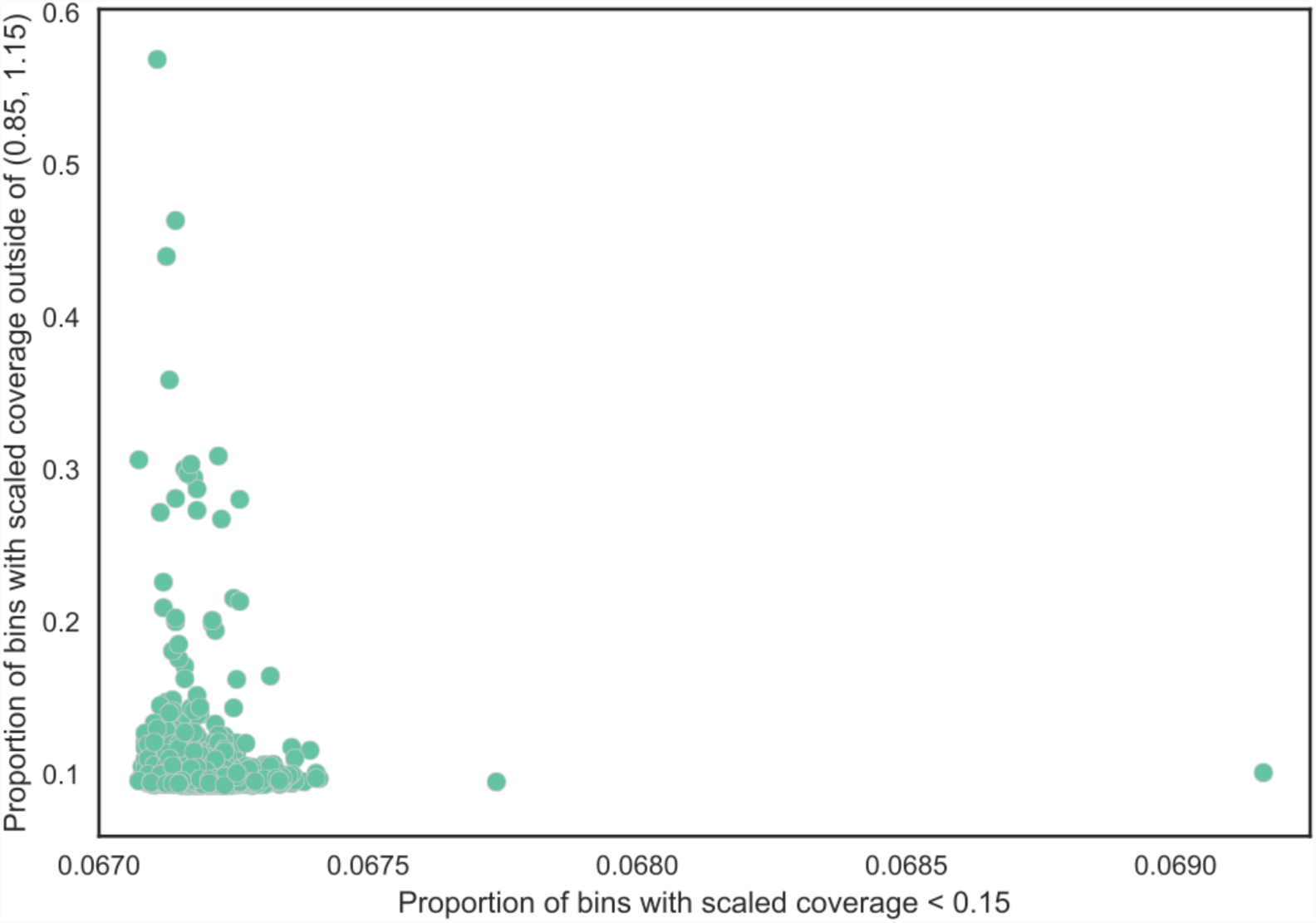
Proportion of 16,384-bp bins where the estimated coverage is less than 0.15 on the x-axis and outside of (0.85, 1.15) on the y-axis among 2,076 human WGS samples. High values on the x-axis indicate large areas with low or no coverage. Values on the y-axis indicate samples with a large bias-with high variance in coverage values.

### Interactive visualization

To facilitate visualization and rapid sample quality control, *indexcov* aggregates all of the output plots into a single, integrated HTML file. Because web browsers struggle to plot highly complex datasets, *indexcov* also generates an overview page that contains static thumbnail images of per-chromosome data. When a user clicks on a thumbnail of interest, she is taken to the full, interactive version of that chromosome plot. In most cases, it will be clear from the thumbnail that there is nothing of interest in that chromosome, so more detailed exploration will not be needed.

The overview page is laid out such that the “sex plot” (e.g., **Figure 3**) and the “tile plot” (e.g., **Figure 4**) are at the top, since these visualizations have the highest information density and are therefore most likely to be immediately useful to the user. If there are major problems with the coverages profiles across an entire cohort, it will be immediately visible from these plots. Subsequently, the PCA plots for batch effects are displayed along with hyperlinks to download the tab-delimited text files reflecting the raw output of *indexcov*. These raw data files include a BGZIP’ed^13^ BED file containing the scaled coverage for each sample for each 16,384 base tile, as well as a pedigree (PED) file that contains each sample, its inferred sex, estimated copy-number for the sex chromosomes, the first five principal components, and the tile statistics described above.

Finally, the overview page displays sample coverage profiles across the genome. For each chromosome, we display a static image of the coverage distribution plot (e.g., **Figure 2B**), as well as a static image of relative depth along the chromosome (e.g., **Figure 2A**). When a user clicks on either static image for a given chromosome, they are taken to the interactive version of that plot so that they can hover to see outliers or features of interest. A live, interactive example of the resulting HTML output if *indexcov* is available at: http://bit.ly/indexcov-example. Each section in the page includes a link to a help document describing the plot type in that section.

### Speed & Scaling

Since *indexcov* must keep each index in memory, memory-use scales linearly with the number of sample files and the reference genome size. On a standard server, *indexcov* completed an analysis of 45 human WGS BAM files at 60X coverage in about 45 seconds. We have run *indexcov* on cohorts as large as 2,076 samples, and *indexcov* users have reported similar performance in analyses of cohorts at least twice this size. We have made attempts to reduce the memory usage as much as possible. For example, we use only 1 byte per scaled coverage value for each sample that we accumulate for cohort-wide PCA. Since we are focused on large deviations, the memory reduction afforded by using a single byte instead of 4 or 8 is worth the loss in precision. For ~2,000 samples, *indexcov* will require about 30 minutes and about 60GB of memory. We present the speed as approximation since it will largely be determined by the I/O speed of the storage disk. In addition, the memory use will also vary depending on the collection characteristics of the go programming language’s memory garbage collector.

### Installation & Invocation

*Indexcov* is available for download from https://github.com/brentp/goleft as a static binary executable for all major platforms. It is extremely simple to use, as its sole inputs are a list of BAM files for the relevant samples (from which it automatically locates the associated indexes), as well as the directory to which the BED and HTML output should be written. An example usage for a set of BAMs would look like:

~~~
goleft indexcov -d output-dir/ inputs/*.bam
~~~

and for CRAMs:

~~~
goleft indexcov -fai $fasta.fai -d output-dir/ inputs/*.crai
~~~

## Conclusion

*Indexcov* enables coverage profiling at low computational cost, provides an interactive output that facilitates the detection of coverage anomalies such as aneuploidies, megabase-scale deletions and duplications, and sex chromosome anomalies. The approach is amenable to whole-genome sequencing datasets from any species, and as such, it represents an important and simple to use quality control step.

## Acknowledgements

We thank the Autism Sequencing Consortium’s Whole Genome Sequencing Working Group for their contributions in processing the WGS data from the Simons Simplex Autism cohort.

## Supplement

**Supplemental Figure 1.**
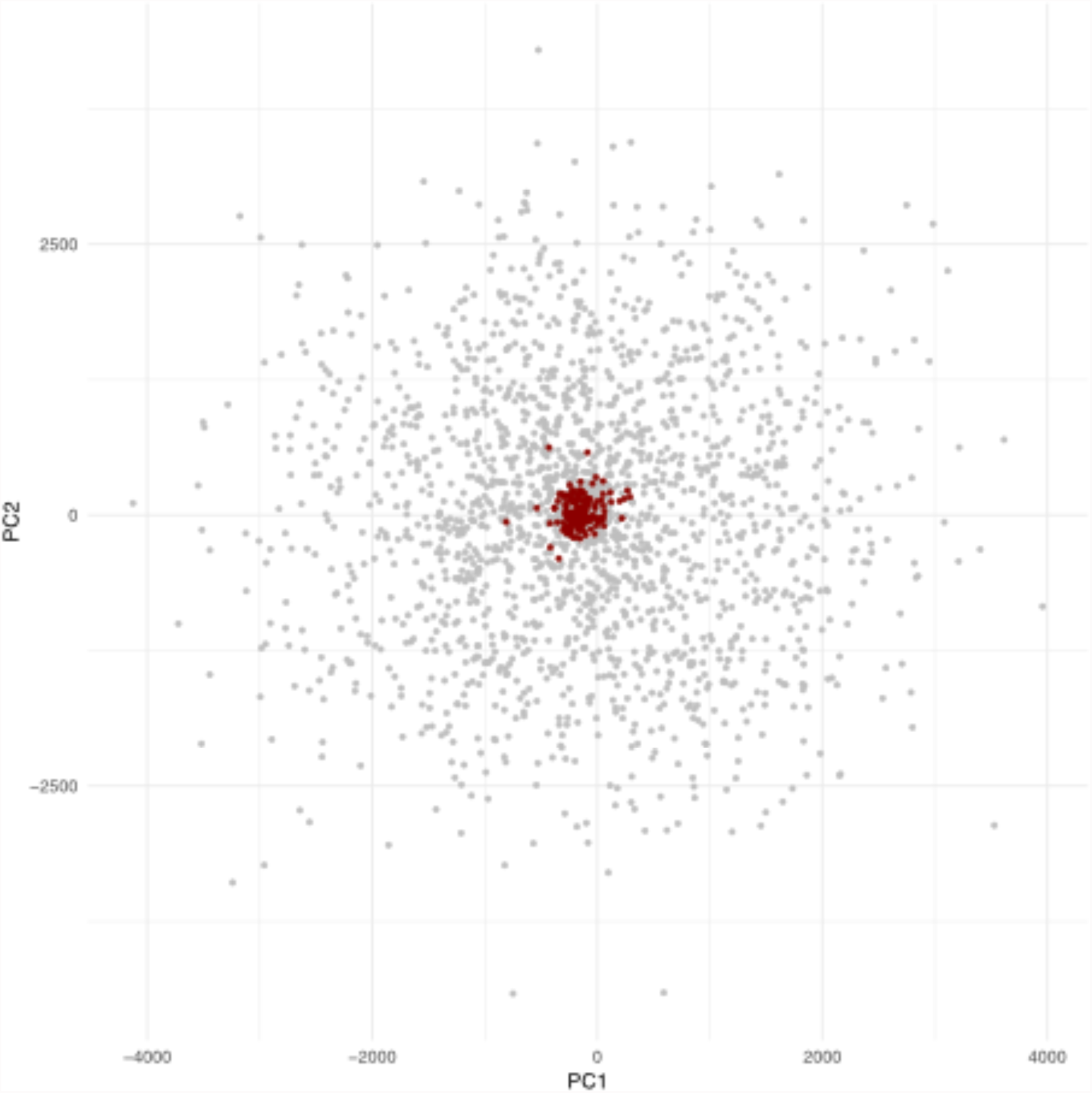
For the 2,076 samples from the Simons Simplex Autism cohort, we plot the first 2 principal components. Samples that were prepared with a PCR-free method are shown in red, with the remaining samples in gray. We see that the samples that had a PCR step have a much greater spread in the principal component values due to the greater variation in genomic coverage resulting from PCR amplification prior to sequencing^9^.

